# Net effects of field and landscape scale habitat on insect and bird damage to sunflowers

**DOI:** 10.1101/804328

**Authors:** Sara M. Kross, Breanna L. Martinico, Ryan P. Bourbour, Jason M. Townsend, Chris McColl, T. Rodd Kelsey

**Affiliations:** Department of Ecology, Evolution and Environmental Biology, Columbia University, 1200 Amsterdam Avenue, New York, NY, USA; Department of Wildlife, Fish and Conservation Biology, University of California, Davis, 1 Shields Ave, Davis, CA, USA; The Nature Conservancy, 555 Capitol Avenue, Ste 1290, Sacramento CA, USA; Hamilton College, 198 College Hill Road, Clinton, NY, USA

**Keywords:** agroecology, crop damage, ecosystem services, farm, hedgerow, integrated pest management, pest control, landscape

## Abstract

Agriculture-dominated landscapes harbor significantly diminished biodiversity, but are also areas in which significant gains in biodiversity can be achieved. Planting or retaining woody vegetation along field margins can provide farmers with valuable ecosystem services while simultaneously benefitting biodiversity. However, when crops are damaged by the biodiversity harbored in such vegetation, farmers are reluctant to incorporate field margin habitat onto their land and may even actively remove such habitats, at cost to both farmers and non-target wildlife. We investigated how damage by both insect pests (sunflower moth, *Homoeosoma electellum*) and avian pests to sunflower (*Helianthus annuus*) seed crops varied as a function of bird abundance and diversity, as well as by landscape-scale habitat. Surveys for insect damage, avian abundance, and bird damage were carried out over two years in 30 different fields on farms in California’s Sacramento Valley. The mean percentage of moth-damaged sunflowers sampled was nearly four times higher in fields that had bare or weedy margins (23.5%) compared to fields with woody vegetation (5.9%) and decreased in both field types as landscape-scale habitat complexity declined. Birds damaged significantly fewer sunflower seeds (2.7%) than insects, and bird damage was not affected by field margin habitat type, landscape-scale habitat variables, or avian abundance, but was significantly higher along field edges compared to ≥ 50m from the field edge. Avian species richness nearly doubled in fields with woody margin habitat compared to fields with bare/weedy margins in both the breeding season and in fall. These results indicate that the benefits of planting or retaining woody vegetation along sunflower field margins could outweigh the ecosystem disservices related to bird damage, while simultaneously increasing the biodiversity value of intensively farmed agricultural landscapes.

## Introduction

In the face of significant losses of both diversity and abundance of avian species (Rosenberg et al. 2019), farming agroecosystems represent a critical frontline for improving vast tracts of land for breeding, migrating, and overwintering birds. Agricultural intensification can drive biodiversity loss, but, paradoxically, agricultural systems rely on the ecosystem services provided by biodiversity (Johnson et al. 2017). Establishing and protecting agroecosystems that take advantage of functional diversity to provide ecosystem services at the farm and landscape level is a way to simultaneously decrease chemical inputs and increase biodiversity (Daily et al. 2000, Weier et al. 2018, Kleijn et al. 2019). To this end, there have been calls for biodiversity conservation to be expanded beyond the reserve system, for example by conserving and promoting functional diversity in expansive agricultural settings (Kremen and Merenlender 2018, Grass et al. 2019). Bringing together these two mindsets can create a win-win situation for both conservation and agriculture. For example, establishing or maintaining strips of woody vegetation along field margins can increase the diversity, abundance, and corresponding ecosystem services, of pollinators (Garibaldi et al. 2011, M’Gonigle et al. 2015, Sardiñas et al. 2016), arthropod predators (Eilers & Klein 2009; Gareau, Letourneau & Shennan 2013), and birds (Heath *et al*. 2017).

Farmers are the primary decision makers for land management choices within agricultural regions, and their decisions are mostly based on direct economic returns (Kleijn et al. 2019). Birds are highly detrimental pests to a number of crops worldwide (De Grazio 1978, Gebhardt et al. 2011, Kross et al. 2012, Schäckermann et al. 2014), although the actual costs of bird foraging on crops are rarely quantified because the timing of bird damage often overlaps with crop harvesting. Farmers that perceive birds as detrimental to their crops will take action to deter birds (Kross et al. 2018), often by removing field margin habitat (Gennet et al. 2013) or utilizing commercially available bird deterrents such as gas guns, reflective tape, or netting (Baldwin et al. 2013), all of which can be costly for both farmers and non-target wildlife. Bird depredation of crops therefore not only has direct economic implications for growers, but can lead farmers to oppose conservation programs within agricultural communities and on their own properties (Kross et al. 2018).

Studies into the detrimental behaviors of birds rarely focus on potentially beneficial impacts, and similarly, studies into the beneficial pest-control services of birds rarely focus on the fact that the same species may cause damage to crops (Pejchar et al. 2018) with a few recent exceptions (Peisley et al. 2016, Gonthier et al. 2019). The effects of natural vegetation on biological control can vary with crop type, seasonality, farm management, and the demographic effects of interactions between natural enemies and pests (Karp et al. 2018, Settele and Settle 2018). Therefore, disentangling the complex relationships between landscape- and field-level habitat complexity and crop damage from insect and avian pests- and communicating these results to farmers and policymakers- has critical implications for habitat management in agroecosystems.

In California, one of the world’s most productive and intensive farming regions, less than 4% of potential field margins have been planted with woody vegetation such as hedgerows (Brodt et al. 2009); field margins therefore have significant potential for increasing the biodiversity conservation value of farmland. However, farmers rank uncertainty around the potential benefits of hedgerows and the possibility that these hedgerows could harbor plant, insect and vertebrate pests as constraints to adopting the practice (Brodt et al. 2009). Research to provide information about the costs and benefits of retaining or planting such habitats is therefore critical to inform land management decisions. Here, we present a study to investigate the effects of field-margin and landscape-scale habitat on insect and bird damage to sunflower (*Helianthus annuus)* crops in California.

## MATERIALS and METHODS

### Study Area and Crop

California’s Central Valley runs 724 kilometers north-south and covers a total of 10.9 million hectares (26.9 million acres). It is one of the most productive agricultural landscapes in the world, producing over 25% of the fresh produce consumed in the United States (USDA 2015), and valued at over $45 billion (USD) per year. Over 95% of the Central Valley’s riparian and wetland ecosystems have been replaced by highly intensive agriculture and urban development (Katibah 1984, Frayer et al. 1989), with remnant native habitat existing only in fragmented and isolated patches. Nevertheless, some native biodiversity in this region persists despite the highly human-modified landscape (Heath et al. 2017).

Each year, sunflower is grown for hybrid seed production on an average of 20,234ha (50,000 acres) across California’s Sacramento Valley, producing over 31,750 tons valued at approximately $70 million/year (Long et al. 2019). California’s Central Valley produces over 95% of the United States’ hybrid sunflower seeds, and over 25% of global sunflower seeds (Long et al. 2019). Sunflowers grown for seed are valued at five to ten times that of the commercial oil crops for which they are used (Long et al. 2019), and growers therefore have a low threshold for damage. All sunflower fields in our study were grown for the same seed company and therefore were grown using the same field-management practices. This study was conducted within conventional fields (i.e. non-organic fields), but no growers reported utilizing insecticides on their fields over the duration of this study.

The predominant insect pest for sunflowers in North America is the sunflower moth (*Homoeosoma electellum*). Female sunflower moths lay eggs among the florets of sunflowers in early bloom, and eggs take 2-5 days to hatch. After hatching, larvae remain on the face of flowers for 8 days before boring into the developing seeds where they can cause losses of 30-60% of a crop (Long et al. 2019). Birds are also a key pest of sunflower crops around the world (De Grazio 1978, Schäckermann et al. 2014, Long et al. 2019, Ernst et al. 2019). Within a field, bird damage to sunflowers is often concentrated to the edges nearest to habitat that can act as shelter for birds. For example, in Israel, bird damage within a field was highest in areas close to trees (>5m in height), but increasing the number of trees within a 1-km radius of fields was not associated with higher damage (Schäckermann et al. 2014), suggesting that presence of habitat along edges of crops prone to bird damage is more important than the presence of habitat in the landscape overall.

### Field- and Landscape-Habitat Complexity

We conducted bird counts and collected sunflower damage data from six fields with woody margin habitat and seven fields with bare or weedy field margins in 2014, and from 12 complex fields and 5 simple fields in 2015, for a total of 30 fields sampled. To quantify local (field) habitat complexity, we collected data on the height, width, and number of canopy layers of field margin vegetation at 5 locations along each transect (see Heath et al. 2017 for details). To quantify and incorporate landscape habitat complexity into our study design, we selected fields at varying distances from natural habitat, which in our study area consists mainly of remnant and restored riparian areas (Figure 1). We used pre-existing habitat data for our study area (CA DWR 2008, Geographic Information Center 2009), and added by hand any trees within 800m of each transect that were not included in the existing dataset (e.g. trees lining driveways, trees around homesteads). To calculate the distance to riparian area, we used ArcGIS 10.1 (ESRI 2010) to create a distance raster that encompassed the entire study area by using the Euclidean distance algorithm. We used the riparian vegetation GIS dataset (habitats classified as native riparian, blue oak woodland, valley foothill riparian, fresh emergent wetland, saline emergent wetland, and valley foothill riparian) as the ‘source’ input for the algorithm and set the output grid cell size to 10 meters. Each field’s transect center point was then buffered by 50 meters, and we calculated the distance from each grid cell within the buffer to the nearest riparian vegetation polygon. The mean distance for all cells within each buffer was calculated as the distance value for each field. We also calculated the mean proportion area consisting of natural habitat (Appendix 1) at concentric buffer distances of 100m, 200m, 400m, and 800m, which have been shown to be relevant scales for riparian bird species in the Central Valley (Seavy et al. 2009).

**Figure 1:**
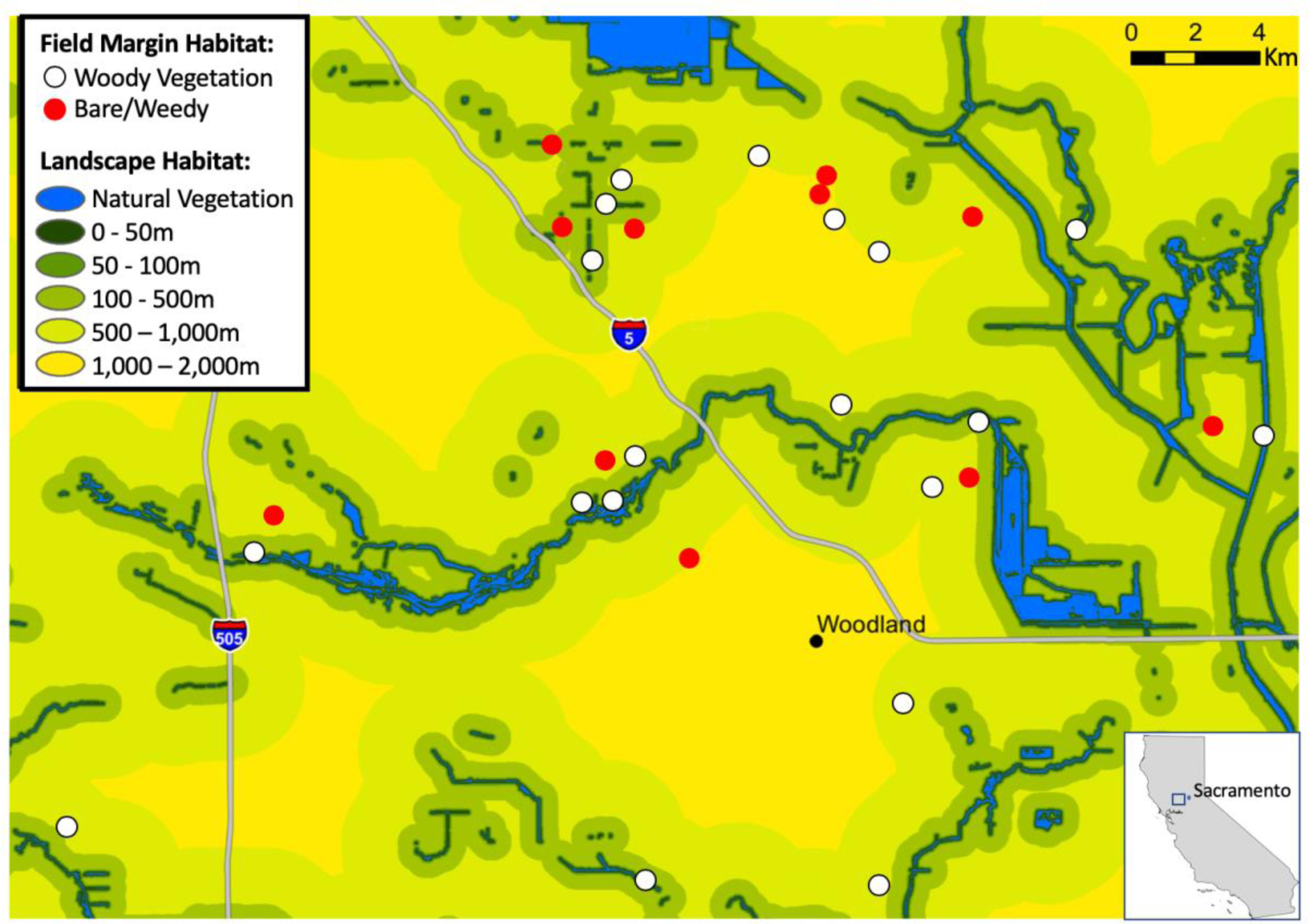
Map showing sunflower field locations at varying distances from natural habitat (blue) across an intensive agriculture landscape around the city of Woodland in the Sacramento Valley of California. Sunflower fields had either bare/weedy field margin habitat (red points), or had woody vegetation field margin habitat (white points).

### Vertebrate Exclosures

In 2015, we created exclosures to prevent vertebrates (birds and bats) from accessing sunflowers (see Maas et al. 2019 for a review of exclosure methods). Enclosures consisted of nylon bird netting (No-Knot Bird Netting ¾” polypropylene mesh, Bird B Gone Inc®, Irvine, CA) draped over an area 4 rows of sunflowers in width and approximately 20 flowers in length and secured to cover the tops of the flowers to a height of approximately 2-4 feet above the ground. Exclosures were installed in late spring, prior to the onset of bloom (which is when sunflower moth typically lay eggs on the flowers), and were checked and maintained over the entire growing season until final damage estimates were made. We set up four exclosures in each field, with the closest end of each exclosure located 5m, 10m, 50m, and 100m from the edge of the field. Due to last minute changes in the harvest schedule at some fields, we were able to collect damage data from the exclosures at nine different fields. All experiments were carried out in accordance with the University of California’s Institutional Animal Care and Use Committee approved protocol #18033

### Sunflower damage

We quantified both bird and insect damage by visually inspecting each of ten sunflowers within each sampling area. Sunflowers were chosen by reaching out to select a plant stalk, so the seed-bearing area of each plant was not seen until after the plant was selected (most sunflowers were at or above head-height for observers). Observers moved a few steps along and between rows to select each new flower. Bird damage was characterized by missing seeds. We were careful to avoid classifying wind-damaged seeds that had been rubbed off of sunflowers by a neighboring flower as bird damage. These seeds were generally removed from larger continuous areas of the sunflower head, whereas seeds removed by birds were in patchy sections or removed singularly. Wind-damaged seeds were also often seen whole on the ground underneath the plants. Insect damage was characterized by an area of visible frass (insect excrement and webbing) on the surface of multiple sunflower seeds. Seeds under the frass were often shrunken or visibly damaged. All areas that were under frass were classified as insect-damaged.

To estimate the percent of seeds on each sunflower that were damaged, we used a pre-cut circular piece of galvanized steel chicken-wire that was marked to allow for easy measurement of the flowers. Sunflower heads were classified into different size classes based on the diameter (to the nearest 1.3 cm, or 0.5 inches) of the seed-bearing area on each plant. We then estimated the number of hexagons on the wire (to the nearest ¼ hexagon) that was damaged by birds or damaged by insects on each sunflower head. Using the flower circumference and the known area within each hexagon of our grid, we were then able to calculate the percent of each sunflower head that was damaged by birds, and the total that was damaged by insects. To estimate yield, damage from insects and damage from birds were summed for a total percent damage to each sunflower, since both types of damage result in a direct loss of yield for growers.

We sampled from 10 sunflowers at distances from 0m to 200m from the field edge. In 2014, we collected observations of both insect and bird damage from each site at 0, 10, 20, 30, 40, 50, 75, 100, 150, and 200m from the field edge. In 2015, we collected observations from each site at 5, 10, 50, and 100m from the field edge because we found in 2014 that bird damage dropped to close to 0 at distances beyond 50m, and that insect damage was largely unchanged by distance from the field edge (see Figure 2). Estimates for insect and bird damage in 2015 were taken from sunflowers within exclosures and from sunflowers that were approximately 10m from the exclosures (parallel to the field margin), but only data from non-enclosed sunflowers was used in our comparative analysis of insect damage.

**Figure 2:**
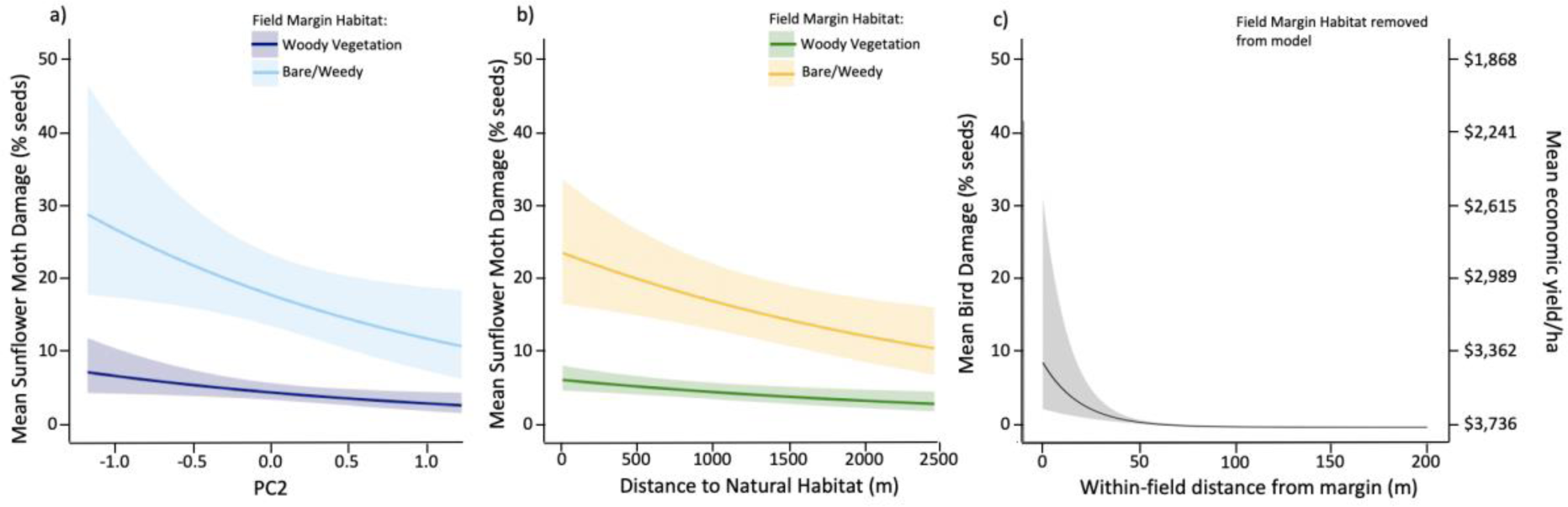
Model estimates of percent of sunflower seeds damaged as a function of the presence (darker colored lines) or absence (lighter colored lines) of woody vegetation along field edges and, a) an orthogonal axis for field margin habitat type, b) the distance to the nearest natural habitats; and c) percent seeds damaged by birds as a function of the distance of sampling points within each field from the nearest field margin. The mean economic yield per hectare is shown as a secondary y-axis and applies to all three panels.

### Bird counts

We conducted four bird surveys at each site, two in summer (June 9 - July 2) and two in fall (August 5- September 16). All bird surveys were conducted by trained observers and timed to coincide with sunflower bloom in the summer (when sunflower moths typically lay eggs on the flowers), and immediately prior to the seed harvest in the fall. All counts were conducted between dawn and 10am and were not conducted in very cold (<3C) or very hot weather (>24C), in high winds or heavy precipitation. Counts were also re-scheduled if there were any farm workers or machinery in our focal field. We conducted two counts per visit at each field: one to quantify the birds utilizing the field margin habitat, and another to quantify the birds utilizing the field interior. These methods provide relative values for comparing inter-site bird communities. To count birds utilizing field margin habitat, observers walked a 200m transect slowly over 10 minutes, counting all detectable birds by sight or sound within 20m of the field margin, but not within the field itself. To count birds utilizing the field interior, observers returned to the mid-point of the transect, allowed five minutes for birds to settle, and then conducted a 10-minute point count focused only on birds that were observed within the field. We counted all birds detected within each field because each species was assumed to have similar detectability in all fields, since sunflowers were at similar levels of maturation and height at the time of each count, and since fields were all of a similar size. We used different methods for the edge and interior transects to maximize our detection of birds utilizing each type of habitat. While these methods may result in counting the same individual in both habitats on the same visit, this is relevant since birds at our study sites were regularly observed using both the field margin and field interior habitats.

### Statistical Analyses

Because the variables describing field margin habitat (height, width, and number of vegetation layers) were highly correlated, we used a Principle Components Analysis (PCA) to reduce these into two orthogonal axes that explained over 95.5% of the variance among them. The two axes, PC1 and PC2, were included as predictor variables in our candidate models for sunflower damage and for bird abundance and richness. PC1 explained 86.2% of the variability among habitat variables and was negatively associated with all three variables, whereas PC2 was positively associated with habitat width and height, and negatively associated with habitat layers. Therefore, if PC1 is a positive predictor of damage, we would expect less damage at sites with habitat that is taller, wider and has more layers (because of the inverse relationship). If PC2 is a positive predictor of damage, we would expect less damage at sites with more habitat layers and more damage at sites with taller/wider habitat. We also found collinearity among the predictor variables for landscape-scale habitat complexity, so constructed separate models for each landscape-scale habitat complexity variable. Model selection revealed that the variable for mean distance to natural habitat was most parsimonious in our sunflower damage models (Tables S1-2), so we present the results from that model in the main text of this paper.

We used a Wilcoxon rank-sum test to compare the total insect damage observed inside exclosures and in adjacent non-exclosure locations. For all other analyses, only the data from the non-exclosure locations were used for investigating the effects of habitat variables on sunflower damage.

For both damage categories, we used generalized linear models with a negative binomial family of errors to analyze our data on percent damage to sunflowers in R v.3.3.1. Sunflower moth damage and bird damage were analyzed in separate models. For our bird abundance and richness data, we ran eight separate linear regressions for avian species richness and abundance along the field edge and within the field interior for data collected in summer and in fall. For all analyses, we included as predictor variables in our maximal models the continuous variables for the distance from the nearest riparian habitat, PC1, and PC2, as well as the categorical variable for whether the field had a weedy or bare edge (simple edge habitat) or had woody field margin habitat (complex edge habitat). We simplified the maximal models by removing interactions, then main effects, until no further reduction in residual deviance (measured using Akaike’s Information Criterion) was obtained. For all regression analyses, we considered candidate models with **Δ**AIC ≤ 2 and chose the most parsimonious model.

### Economic Estimates

We used published data on the range and mean sunflower yields and economic value for the Sacramento Valley from 2015-2018 (Long et al. 2018) to calculate the reduction in gross earnings for farmers as a result of insect and bird damage in response to significant predictor variables. Mean sunflower yields were 1,260 lbs/acre (1,412kg/ha; range 1,076-1,748 kg/ha) after seed companies clean and remove nonviable seeds and non-seed material from field harvests (Long et al. 2018). Seeds were valued at a mean value of $1.2/lb ($0.54/kg; range of $0.41-0.68/kg (Long et al. 2018)). We calculated the economic effect size of insect or bird damage by multiplying the scaled effect sizes from our model estimates.

## RESULTS

### Vertebrate Exclosures

There was no significant difference between sunflower damage from insects inside exclosures (mean= 3.40 ± 0.61% damage) compared to areas outside of exclosures that birds and bats could access (mean= 3.08 ± 0.47% damage).

### Sunflower damage

Sunflower moth damage was almost four-times higher at sites with bare or weedy field margin habitat (23.46 ± 1.41%) compared to sites with woody vegetation along field margin habitat (5.89 ± 1.16%; *z=* 7.12, *p* < 0.001). There was a slight decrease in sunflower moth damage as PC2 increased (*z=* -2.75, *p* = 0.005; Figure 2a), and a significant reduction in damage as distance from natural habitat increased (*z=* -2.25, *p* = 0.02; Figure 2b). Bird damage was highest at the edge of fields, regardless of the presence of field margin habitat, and dropped quickly to near 0% within 50m of the field edge (Figure 2c). This effect was driven entirely by distance from field edge, with only the linear (*z=* -4.45, *p* < 0.001) and quadratic values (*z=* 2.98, *p* = 0.003) for distance from field edge retained in the final model.

### Economic Estimates

Our models estimate that at sites adjacent to natural vegetation, farmers would expect to lose $877/ha in lost yields due to sunflower moth damage at sites with bare/weedy vegetation along the field margin, compared to $220/ha in lost yields due to sunflower moth damage at sites with woody vegetation. To put this into perspective, the mean cost of applying insecticides to treat for sunflower moth is $292/ha, so our results suggest that fields in this scenario would be likely to remain under an economic threshold to trigger growers to apply insecticides. In the same scenario, bird damage at the field edge would result in $100 in lost yields but that would decline to negligible damage within 50m of the field edge.

### Bird results

Species richness of complex fields was higher in fields with woody margins in both summer (19.70 ± 0.91) and fall (16.0 ± 1.04) compared to fields with bare/weedy margins in summer (10.4 ± 0.96) and fall (8.17 ± 0.67). We observed 70 different avian species during our summer counts, and 74 species during our fall counts. These included California ‘Bird Species of Special Concern’ (Shuford & Gardali 2008) like northern harrier (*Circus hudsonius*), yellow warbler (*Setophaga petechia),* and California ‘Threatened’ species like Swainson’s hawk (*Buteo swainsoni*), and tri-colored blackbird (*Agelaius tricolor*, 13 individuals observed at one site).

During our summer counts, 64 different bird species utilized sunflower field edges and 49 species utilized field interiors. During our fall counts, we observed 69 species utilizing sunflower field edges and 46 species utilizing field interiors. Further details of bird species observed can be found in Bourbour et al. (In prep).

For our summer counts, avian species richness (t = -5.44, *p* <0.001, Figure 3a) and abundance (t = -5.47, *p* <0.001, Figure 3b) along field edges had a strong negative correlation with PC1. Since PC1 was negatively associated with all three measures of field margin habitat complexity (habitat height, width, and number of canopy layers), our results predict that as field margin habitat becomes more complex, avian richness and abundance along field edges increased. For summer field interiors, avian species richness was uncorrelated with PC1 (t = - 1.83, *p* = 0.08, Figure 3c). None of our predictor variables were retained in the model for avian abundance within the field interior in summer.

**Figure 3:**
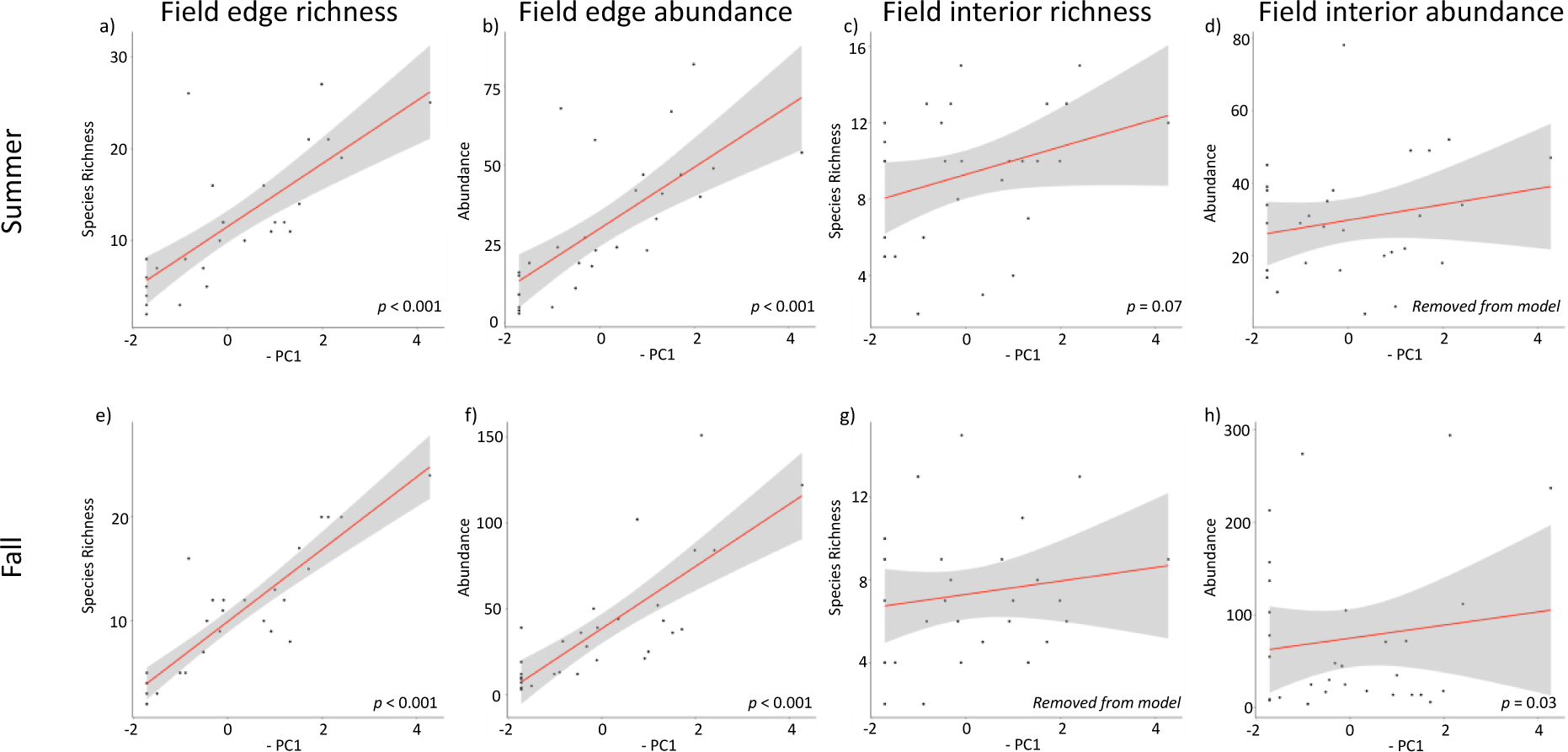
Avian species richness and abundance along sunflower field edges and within sunflower field interiors in Summer (top row) and Fall (bottom row) as a function of increasing field margin habitat height, width, and number of canopy layers (- PC1). Statistical significance of PC1 variable as a predictor in a linear regression for each response variable is shown bottom right in each panel.

In the fall, field edge richness, field edge abundance, and field interior abundance were all positively associated with increasing field margin habitat complexity. Field edge avian species richness was negatively associated with PC1 (t = -9.82, *p* <0.001, Figure 3e), PC2 (t = - 2.80, *p* <0.01) and average distance to nearest riparian habitat (t = -2.30, *p* = 0.03). Avian abundance at the field edge in fall was negatively associated with PC1 (t = -23.40, *p* <0.001, Figure 3f). Avian species richness in field interiors during the fall was not correlated with edge complexity or distance to riparian habitat. Avian abundance was significantly higher at sites with weedy/bare edges, compared to sites with woody vegetation (mean of 109 more birds at simple sites; t = 2.33, *p =* 0.03), but increased in both bare/weedy and woody vegetation field margin types with increasing field margin habitat complexity (negatively association with PC1; t = -2.31, *p* = 0.03, Figure 3h).

## DISCUSSION

Our results suggest that sunflower growers would benefit from planting or maintaining woody vegetation alongside their fields since sunflower moth damage was significantly higher at sites without field margin vegetation, while bird damage was not driven by field margin habitat. Furthermore, within sunflower fields across all distances from the field margin, sunflower moth damage was significantly higher than bird damage, and was therefore the main source of yield loss for sunflower growers in our area. The pest control service benefits that farmers receive from field margin vegetation therefore outweigh the potential ecological disservices associated with bird damage to sunflowers. In fact, bird damage at our 30 fields was similar across sites with and without field margin habitat. Our results also indicate a clear benefit for biodiversity, with significantly higher species richness and avian abundance along field edges that had woody habitat. Combined, these results support the assertion that diversified farming systems can provide both farmers and broader society with multiple additive ecosystem services (Kremen & Miles 2012).

Our exclosures did not reveal an effect of bird foraging on sunflower moth damage. This could be the result of small sample size (n=36 exclosures in 9 fields), or these results could indicate that foliage-gleaning birds and bats are not a major predator of sunflower moth. We predict that the patterns of sunflower moth damage we observed were driven by either increased predation pressure from invertebrates or from aerially-hunting bats and birds. If adult moths are depredated in flight, prior to ovidepositing on flowers, our exclosures would not have detected an effect of aerially-hunting vertebrate predators such as bats and birds.

Whereas if most depredation occurs to adult moths as they lay eggs, to the eggs themselves, or to larvae, our exclosures would have indicated if vertebrate predators were the cause. Because of their nocturnal nature, sunflower moth adults are likely to be targeted more by nocturnal arthropod predators and/or bats (both of which would not be affected by the presence of exclosures) than by the predominantly diurnal avian predators. There is also a possibility that the presence of woody vegetation creates a physical barrier to fields or that sunflower moths avoid areas near woody vegetation because of the potential for higher predation. Further research is clearly needed in this system.

The value of insect pest control provided to US farmers by beneficial insects was estimated at $4.5 billion per year in 2006 (Losey and Vaughan 2006), and the value of insect pest control provided to farmers of corn globally was estimated to be worth over $1 billion per year in 2015 (Maine and Boyles 2015). Studies in California have shown that the presence of habitat along field margins is associated with increased diversity and abundance of beneficial insects including natural enemies (Eilers and Klein 2009, Gareau et al. 2013, Morandin et al. 2014), and with increased bat activity (Kelly et al. 2016). For example, in almond orchards, higher proportions of natural habitat surrounding orchards resulted in higher parasitoid and vertebrate control of the naval orangeworm (Eilers and Klein 2009, Morandin et al. 2014).

Our model selection process revealed that the distance to nearest natural vegetation was the strongest of our landscape-scale predictors of sunflower damage (Table S2). This variable has been shown to be a significant predictor of bird use of agricultural fields in our study region (Kross et al. 2016, Heath et al. 2017) since the landscape is largely dominated by farm fields, with widely spaced corridors of natural habitat along riparian areas, and farmsteads acting as small islands of natural habitat (Figure 1). In this landscape, hedgerows themselves are an important sources of pollination services (Sardiñas et al. 2016) and for supporting pollinator metacommunity dynamics (Ponisio et al. 2019)

The benefits and costs of bird presence on farms are complicated (Pejchar et al. 2018). Individual species can be beneficial to a crop in some seasons and detrimental in others, or may benefit one crop and cause damage to another. Birds may also disrupt other natural trophic cascades that benefit farmers (Grass et al. 2017). Seasonality of avian foraging guilds is important, and often overlooked by either those interested in describing only pest-control services or those interested in describing only damage from pest bird species. For example, red-winged blackbirds (*Agelaius phoeniceus*) are a notorious pest of sweet corn crops in the North American Midwest in late summer and early fall when corn kernels are ripening, but these abundant birds also consume large quantities of a number of insect pests of corn in spring and early summer (Dolbeer 1990; Tremblay, Mineau & Stewart 2001). In sunflowers, some species considered to be major pests of sunflower crops, including blackbird species (Icteridae) and European starlings (*Sturnus vulgaris*), are insectivorous during the breeding season - the time of year that sunflowers are attacked by insect pests, including sunflower head moth. These species later become pests when they switch to a primarily granivorous diet in the fall. Farmers may therefore want to retain, or even encourage, blackbird populations on their land in spring and summer, and then utilize bird deterrent techniques, alternative food sources, and bird-resistant cultivars to reduce damage from birds once crops become susceptible. Importantly, while our results indicate a net benefit of hedgerows for both sunflower yields and avian diversity in California, sunflowers in other regions (Schäckermann et al. 2014, Ernst et al. 2019) suffer from economically significant bird-damage to the same crop. Therefore, we caution that land managers and scientists should consider local climate, habitat availability, agricultural practices, and avian communities before translating our findings into management changes in other regions.

## CONCLUSIONS

Contrary to common assumptions about avian pests, we found that sunflower fields with woody vegetation along their margins did not suffer from significantly higher bird damage compared with fields that had weedy or bare edges. Instead, overall sunflower seed yield was driven by insect damage, which was lower in fields with vegetated margins. Our results show that planting or retaining woody vegetation along field margins can simultaneously decrease insect pest damage to crops (Figure 2) and increase the biodiversity value of sunflower fields for birds (Figure 3), adding to the growing body of scientific studies that demonstrate the benefits of planting or retaining habitat for wildlife along field margins. These results are particularly important for avian conservation in intensive agricultural landscapes, where little natural habitat remains and significant gains in habitat may be made through restoration activities along the ∼96% of field margins in California that are currently bare. Our results demonstrate one case where increasing the habitat value of non-production areas in intensive, conventional farming systems may simultaneously increase yield and benefit biodiversity.

## AUTHOR’S CONTRIBUTIONS

SMK, TRK and JMT conceived the ideas and designed methodology; SMK, BLM, RPB and field assistants collected the data; CM performed the landscape analysis; SMK analyzed the data; SMK led the writing of the manuscript. All authors contributed critically to the drafts and gave final approval for publication.

## ACKNOWLEDGEMENTS

We thank the landowners and growers who provided access and information for this study especially Button & Turkovich, Joe Muller & Sons, Bullseye Farms, Citrona Farms, and Bypass Farms. Pioneer Hi-Bred International allowed for this research to be conducted, and we received significant logistical advice from A. Anderson. We also received invaluable study design advice from R. Long and numerous local pest control advisors. K. Shaw, S. Lei, and E. Barry helped with field work. Fieldwork was funded by the David H. Smith Conservation Research Fellowship to SMK, who was hosted by J. Eadie at UC Davis.

## DATA ACCESSIBILITY

Data will be made available from the Columbia University Library Digital Repository.

### Online Appendices

Supplementary Table 1: Model selection for candidate models explaining sunflower moth damage to sunflower seeds using the distance to nearest natural habitat as a measure of landscape-scale habitat complexity.

Supplementary Table 2: Model selection for candidate models explaining sunflower moth damage to sunflower seeds using, as a measure of landscape-scale habitat complexity, the proportion of natural habitat within concentric distance buffers from each site.

## Supplementary Material

**Supplementary Table 1:** Model selection for candidate models explaining sunflower moth damage to sunflower seeds using the distance to nearest natural habitat as a measure of landscape-scale habitat complexity. A principal components analysis was used to consolidate local habitat complexity variables into two orthogonal axes (PC1 and PC2). Field margins for each site were categorically defined based on the presence or absence of woody vegetation along the field margin. The ‘Distance into Field’ measure is the number of meters within the field for each sampling location from the nearest field edge.

| Name | Residual df | Residual Deviance | $\Delta$ AIC | $w_i$ |
| --- | --- | --- | --- | --- |
| Field Margin + Distance to Natural + PC2 | 190 | 218 | 0 | 0.48 |
| Field Margin + Distance to Natural + PC1 + PC2 | 189 | 218 | 1.7 | 0.2 |
| Field Margin + Distance to Natural + PC1 | 190 | 219 | 3.1 | 0.1 |
| Field Margin + Distance to Natural | 191 | 219 | 3.2 | 0.09 |
| Field Margin + Distance to Natural + Distance into field + PC1 + PC2 | 188 | 218 | 3.3 | 0.09 |
| Field Margin * Distance to Natural + Distance into field + PC1 + PC2 | 187 | 218 | 5.3 | 0.03 |
| Null | 193 | 222 | 42.4 | 0 |

**Supplementary Table 2:** Model selection for candidate models explaining sunflower moth damage to sunflower seeds using, as a measure of landscape-scale habitat complexity, the proportion of natural habitat within concentric distance buffers from each site. Only the landscape-scale habitat variable is shown changed in this table based on the most parsimonious model above. A principal components analysis was used to consolidate local habitat complexity variables into two orthogonal axes (PC1 and PC2). Field margins for each site were categorically defined based on the presence or absence of woody vegetation along the field margin.

| Model Terms | Residual df | Residual<br>Deviance | $\Delta$ AIC | $w_i$ |
| --- | --- | --- | --- | --- |
| Field Margin + Distance to Natural + PC2 | 190 | 218 | 0 | 0.59 |
| Field Margin + Prop. Natural 800m + PC2 | 190 | 219 | 1.9 | 0.23 |
| Field Margin + Prop. Natural 200m + PC2 | 190 | 219 | 4.8 | 0.05 |
| Field Margin + Prop. Natural 100m + PC2 | 190 | 219 | 5 | 0.05 |
| Field Margin + Prop. Natural 50m + PC2 | 190 | 219 | 5.4 | 0.04 |
| Field Margin + Prop. Natural 400m + PC2 | 190 | 219 | 5.5 | 0.04 |
| Null | 193 | 222 | 42.4 | 0 |

